# A mobile circular DNA element drives memory-related processes in mice

**DOI:** 10.1101/2025.09.17.676936

**Authors:** Haobin Ren, Hao Gong, Wei-Siang Liau, Qiongyi Zhao, Alexander P. Walsh, Mason R.B. Musgrove, Joshua W.A. Davies, Esmi L. Zajackowski, Sachithrani U. Madugalle, Paul R. Marshall, Tian Lan, Jichun Shi, Jiazhi Jiang, Wei Wei, Xiang Li, Timothy W. Bredy

## Abstract

Extrachromosomal circular DNAs (ecDNAs) are double stranded, closed loop DNA molecules that act independently of the genome. Here we report the discovery of a brain-enriched ecDNA, *ecCldn34d*, that regulates the transcriptional state of plasticity-related genes and is temporally associated with the stability of fear memories. ecDNA therefore represents a novel class of mobile DNA elements that are involved in experience-dependent neuroadaptation.

---

In mice, activity-induced gene expression in the medial prefrontal cortex (mPFC) is required for the formation of fear memory^1,2^. We, and others, have previously shown that various epigenetic mechanisms, including DNA modification and experience-dependent variations in DNA structure states, are critically involved in this process^3,4,5,6^. The realisation that the genome is dynamic and highly responsive to environmental cues has been a major advance in neuroscience; nonetheless, a complete understanding of the molecular mechanisms of brain adaptation remain to be elucidated.

In 1965, Bassel and Hotta observed double-stranded circular DNA molecules independent of the autosomal genome, which they called extrachromosomal circular DNA (ecDNA)^7^. Varying widely in size and abundance, ecDNAs have since been implicated in a multitude of biological processes including gene amplification, chromatin regulation, and structural remodelling of the chromosome^8–11^. Although ecDNAs have been identified in different cell types, whether they are functionally active in the brain is not known.

To begin to address this question, we used a circular DNA sequencing approach to identify ecDNAs in the mPFC of adult male C57Bl6 mice (Fig S1A-C). Although there was substantial individual variability in the representation of ecDNAs across biological replicates, the majority of brain-derived ecDNAs fell within a size range of 100 to 1000 base pairs (Fig. 1A). In addition, the genomic regions from which ecDNAs originate are broadly distributed, suggesting that ecDNAs form stochastically during early brain development (Fig. 1B). An examination of the genomic architecture ecDNA biogenesis revealed that the majority of brain-enriched ecDNAs are derived from intergenic and intronic regions, with only a small fraction harbouring protein-coding sequence (Fig. 1C). In addition, approximately 67% contained repetitive elements, with short interspersed nuclear elements (SINEs) being the most prevalent (30%) (Fig. 1D), indicating that most ecDNAs in the brain may have evolved from non-autonomous retrotransposons.

**Figure 1.**
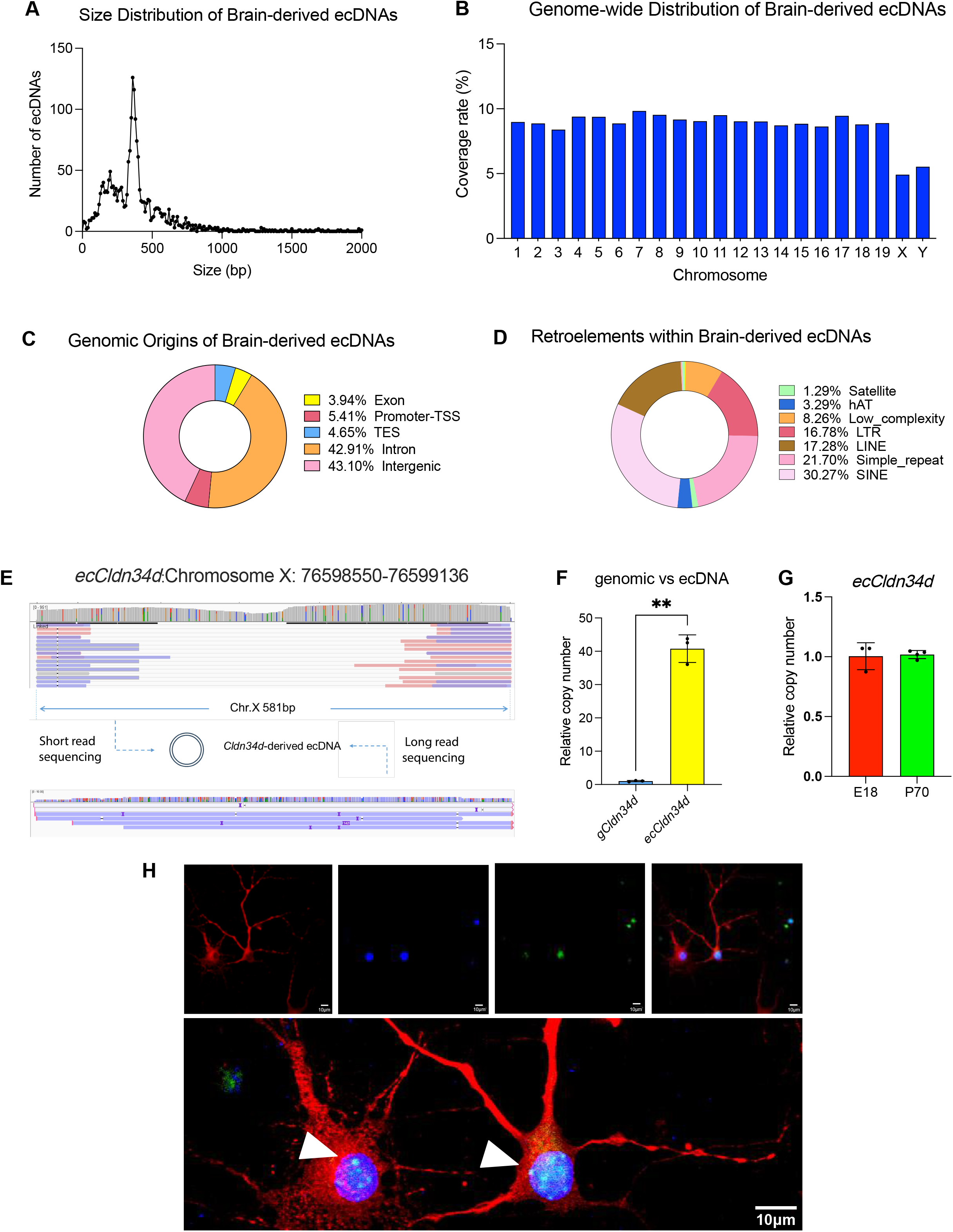
A. Size distribution of neuronal ecDNAs. B. Sequencing coverage of neuronal ecDNAs mapped across each chromosome. C. Annotation of genomic features within ecDNAs identified by Circle-Map. D. Classification of retrotransposon elements that overlap with ecDNA coordinates. E. Representative genome browser view of the *Cldn34d* locus, illustrating *ecCldn34d* detection by both short-read and long-read sequencing. F. Relative copy number of *ecCldn34d* compared to the genomic *Cldn34d* locus in neurons (n = 3 biologically independent samples; unpaired *t*-test; \*\**p* <0.01). G. Relative copy number of *ecCldn34d* in cortical neurons from embryonic day 18 (E18) embryos versus mPFC from postnatal day 70 (P70) mice (n = 4 biologically independent samples; unpaired t-test; *p* = 0.9944). H, Representative DNA FISH image showing *ecCldn34d* signals (indicated by white arrowhead) within the nucleus of primary cortical neurons. Scale bar, 10 µm.

To confirm the existence of brain-enriched ecDNAs, we next performed long-read sequencing on an independent sample of ecDNAs derived from the adult mPFC. In this experiment, ecDNA was isolated selectively from neurons following fluorescence-activated cell sorting with NeuN as a neuronal marker. Focusing on the top 2 million intact reads, we identified 33,227 ecDNAs with breakpoints confirmed using the alignment-based computational method, ecDetector, and an orthogonal informatics pipeline, FLED (Fig. S1A). The median size interval of full-length neuronal ecDNAs was 2373bp, whereas those found in non-neuronal cells were 298 bp (Fig. S1B and C). Similar to the short read data, ecDNAs in neuronal and non-neuronal subcellular populations were more likely to be derived from intergenic and intronic regions (Fig S1D and E). A gene ontology (GO) analysis on genomic loci from which ecDNAs were derived revealed distinct patterns exhibited in neurons versus non-neuronal cells. Notably, genes associated with synapse organization or function were enriched in the NeuN+ group whereas genes involved in nervous system development and cell morphogenesis were enriched in the NeuN-population (Fig. S1F and G).

The circular structure of several ecDNAs that were identified by both the short and long read sequencing was validated using divergent PCR (Fig. S2A). Amongst the top candidates, an ecDNA originating from the first intron of the Claudin 34d (*Cldn34d*) gene locus (Fig. 1E) was selected for further investigation. Following confirmation by Sanger sequencing (Fig. S2B), we noted that *ecCldn34d* contains multiple single-nucleotide polymorphisms, which suggests that it may be potentially modified after its formation (Fig. S2C). Nonetheless, quantitative PCR revealed that there are ~50 copies of *ecCldn34d* per cell and this appears to be stable across the lifespan (Fig. 1F and G). Using probes directed toward the specific back-junction site, we found that ecCldn34d is primarily localized to the nucleus of cortical neurons, a finding which was confirmed following subcellular fractionation (Fig. 1H, and S2D). Together, these data suggest that *ecCldn34d* is a unique and long-lived feature of the neuronal genome, which positions it as a putative candidate involved in transcription or other processes in the nucleus.

An *in vitro* model of activity-dependent plasticity was next used to determine whether *ecCldn34d* dynamically interacts with the linear genome to regulate transcription. dCas9-based *ecCldn34d* immunoprecipitation (DIP) followed by DNA short-read sequencing (Fig. S3A and B) revealed 41,288 peaks in the naive group and 25,944 peaks in activated neurons. Of these, 1,643 peaks were identified enriched upon KCl-induced depolarization, while 17,585 peaks showing higher enrichment in the naive state. Generally speaking, most peaks were located within intergenic and intronic regions (Fig. 2A).

**Figure 2.**
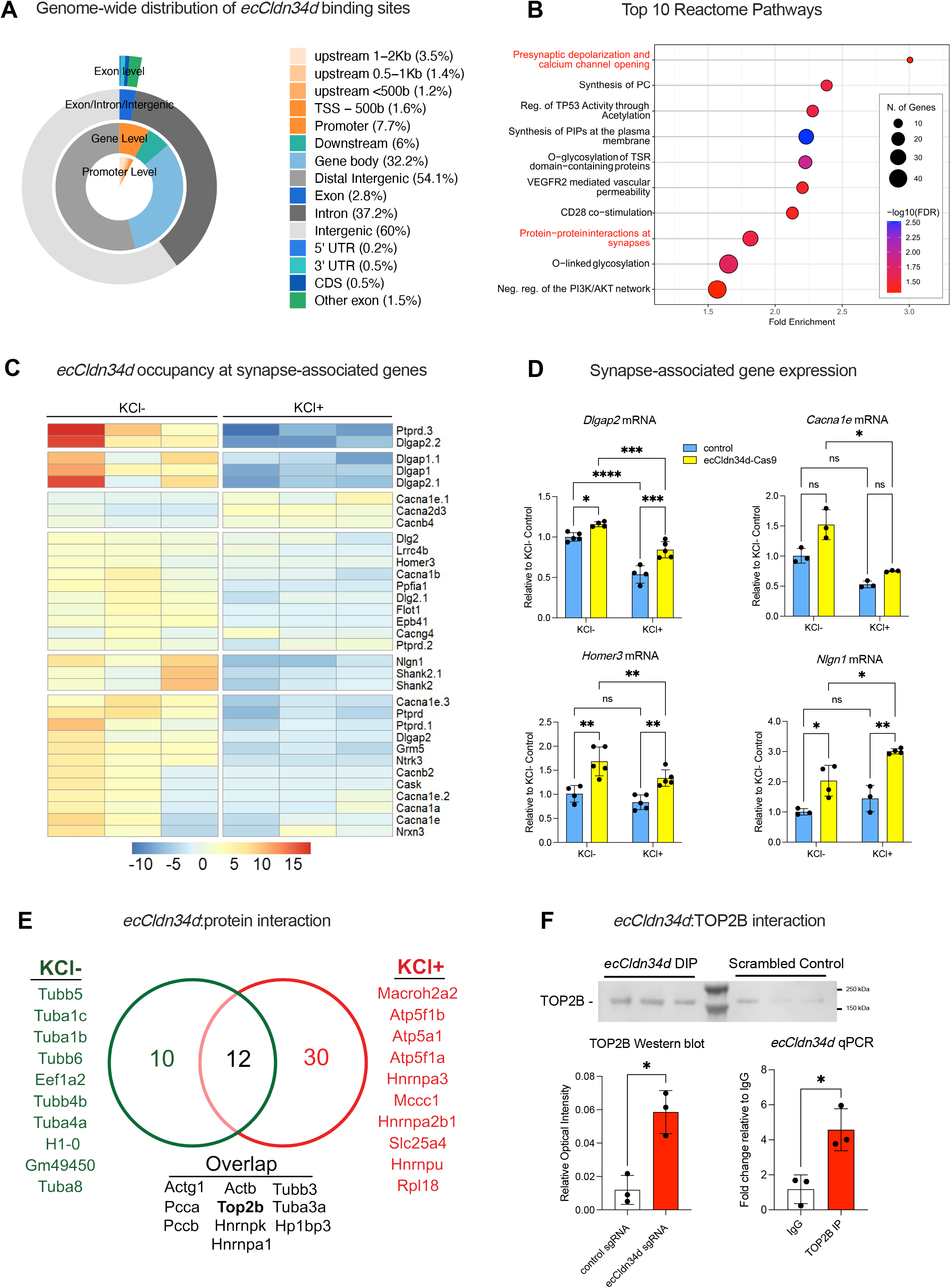
A. Annotation of genomic features within activity-dependent *ecCldn34d* binding sites in neurons. B. Major reactome pathways of genes associated with activity-dependent *ecCldn34d* binding sites. Neuron-related pathways are highlighted in red. C. Heatmap showing *ecCldn34d* occupancy at major synapse-associated genes following neuronal depolarization. D. Relative expression level of typical synapse-associated genes estimated following depolarization in control versus *ecCldn34d* KO neurons (n ≥ 3 biological replicates for each group; Two-way ANOVA analysis; effect of *ecCldn34d* knockout: \*\**p* = 0.0037 for *Dlgap2*, \**p* = 0.0406 for *Cacna1e*, \*\**p* = 0.0067 for *Homer3*, \*\**p* = 0.0064 for *Nlgn1*;effect of neuronal depolarization: \*\*\**p* = 0.0007 for *Dlgap2*, \*\**p* = 0.0067 for *Cacna1e*, \**p* = 0.0229 for *Homer3*, \**p* = 0.0318 for *Nlgn*; For pairwise comparison: \*\*\*\**p* < 0.0001, \*\*\**p* < 0.001, \*\**p* < 0.01, \**p* < 0.05). E, Proteomic identification of *ecCldn34d*-interacting protein partners in neurons upon depolarization; the top 10 proteins are shown. F, Validation of the interaction between TOP2B and *ecCldn34d. Top*, representative western blot for TOP2B after *ecCldn34d* DIP. *Bottom left*, quantification of TOP2B signal intensity from DIP experiments (n = 3 biological independent samples; paired t-test; \**p* = 0.0365). *Bottom right*, qPCR analysis of *ecCldn34d* enrichment in TOP2B immunoprecipitation samples (n = 3 per group; paired t-test; \**p* = 0.0376).

Focusing on gene targets that exhibited the most dynamic changes in *ecCldn34d* occupancy in response to neural activity, a reactome analysis revealed plasticity-related pathways, including presynaptic depolarisation and calcium channel opening and protein-protein interactions at synapses within the top 10 pathways, which indicates a putative role for *ecCldn34d* in regulating key aspects of gene expression associated with neuronal function (Fig. 2B). Upon closer examination of *ecCldn34d* binding at synapse-associated genes, a dramatic reduction in *ecCldn34d* occupancy following KCl-induced depolarisation was observed (Fig 2C), which led us to hypothesise that *ecCldn34d* may broadly serve as a negative regulator of experience-dependent gene expression.

In agreement with this, the expression of several genes whose protein products are directly involved in experience-dependent plasticity was upregulated following *ecCldn34d* knockout in primary cortical neurons, regardless of activation state (Fig 2D and S3C). This includes *Dlgap2* (discs large associated protein 2), a synaptic scaffold protein associated with neurological disorders characterised by memory deficits^12^; *Cacna1e* (calcium voltage-gated channel subunit alpha1 E), an ion channel (CaV2.3) implicated in PTSD^13^; *Homer3* (homer scaffold protein 3), a postsynaptic density protein involved with dendritic spine formation and synaptic tagging^14^; and *Nlgn1* (Neuroligin 1), a memory related postsynaptic adhesion molecule^15^.

Next, to identify potential *ecCldn34d*-protein binding partners that may mediate its effect on gene expression, a dCas9-mediated *ecCldn34d* pulldown followed by mass spectrometry analysis was performed on cross-linked samples derived from activated primary cortical neurons, *in vitro*. Overall, we identified 52 *ecCldn34d*-bound proteins (Fig. 2E), with the majority of chromatin-associated proteins bound to *ecCldn34* following KCl-induced depolarisation. Interestingly, TOP2B (Topoisomerase II Beta), an enzyme that regulates DNA topology and is critically involved in transcription^16^, was bound to *ecCldn34d* irrespective of activation state, which suggests a stable interaction. This interaction was confirmed using reverse Co-IP assays (Fig. 2F).

Finally, to test the hypothesis that *ecCldn34d* regulates fear-related memory, a viral-mediated CRISPR-Cas9 approach was used to reduce *ecCldn34d* copy number in the mPFC either before or after exposure to a cued fear conditioning paradigm (Fig 3A). The *in vivo* efficiency of the Cas9 construct was verified by qPCR, which showed a significant decrease in the copy number of *ecCldn34d* with no change in the *Cldn34d* host gene locus (Fig. 3B and C). *ecCldn34d* knockout one week prior to training (Fig 3D) had no effect on within-session fear acquisition (Fig 3E) or the expression of fear memory when assessed 24 hours post-conditioning (Fig. 3F). However, when the mice were tested 7 days later, there was a highly significant impairment in memory recall (Fig. 3G). In contrast, *ecCldn34d* knockout one day after training (Fig 3H) led to enhanced recall when tested 7 and 14 days after training (Fig 3J and K). Collectively, these findings indicate a temporally relevant role for *ecCldn34d* in the stabilization of long-term fear memories, the precise mechanism of which remains to be determined.

**Figure 3.**
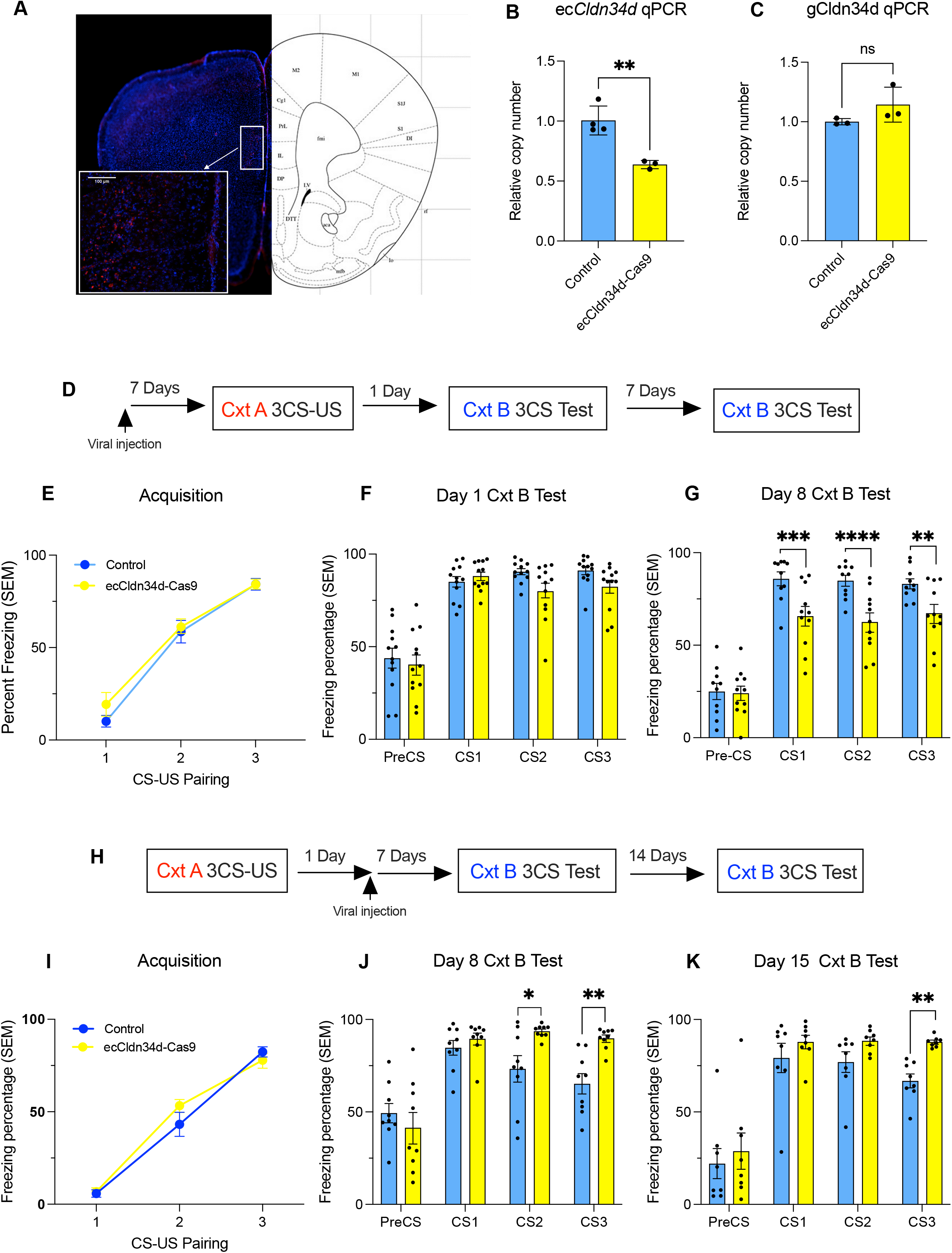
A. Representative immunohistochemical image confirming lentiviral delivery to the mPFC (Scale bar: 100 µm). B-C. qPCR quantification of *ecCldn34d* (b) and the genomic *Cldn34d* locus (c) in the mPFC following transduction with a control or ecCldn34d KO lentivirus (n ≥ 3 biological replicates for each group; unpaired t-test; \*\**p* = 0.0060) D-G, Effect of ecCldn34d knockdown prior to fear conditioning. D. Schematic of the behavioural paradigm. E-G, Behavioural assessment of fear responses in mice receiving control or knockdown virus tested one day (F) and 8 days (G) after fear conditioning (n ≥ 8 biological replicates; Two-way ANOVA analysis; For pairwise comparison: \*\*\*\**p* < 0.0001, \*\*\**p* < 0.001, \*\**p* < 0.01). H-K, Effect of *ecCldn34d* knockdown after fear conditioning. H. Schematic of the behavioural paradigm. I-K, Behavioural assessment of fear responses in mice receiving control or knockdown virus tested 8 days (J) and 15 days (K) after fear conditioning (n ≥ 8 biological replicates; Two-way ANOVA analysis; For pairwise comparison: \*\**p* < 0.01, \**p* < 0.05).

In summary, we have discovered that *ecCldn34d* serves to control neuronal gene expression programs associated with synaptic plasticity, and that it is directly involved in regulating temporal aspects of memory. Mobile circular DNA elements therefore represent a new layer of epigenetic control over experience-dependent neuroadaptation.

## Methods

### Animals

Adult male C57BL/6J mice (9–10 weeks old, weighing 20–25 g) were used in all experiments. Mice were group-housed (four per cage) under a 12-hour light/dark cycle (lights on at 07:00 h) in a temperature-(22 ± 2 °C) and humidity-controlled (50 ± 10%) vivarium, with *ad libitum* access to standard rodent chow and water. Mice were euthanized by cervical dislocation, and the mPFC was rapidly dissected from whole brains and immediately processed for subsequent ecDNA enrichment. All procedures were performed in accordance with protocols approved by the University of Queensland Animal Ethics Committee.

### Fluorescence-activated cell sorting (FACS)

The dissected mPFC was dissociated into a single-cell suspension using FACS lysis buffer (final concentrations: 0.32 M sucrose, 10 mM Tris-HCl, pH 8.0, 5 mM CaCl2, 3 mM MgOAc, 0.1 mM EDTA, 1 mM DTT, 0.3% Triton-X100 and 100× Protease Inhibitor Cocktail) into a single-cell suspension. Suspended cells were then fixed with 1% formaldehyde for 5 min, and the reaction was quenched by the addition of glycine (0.125 M final concentration). Following two washes with ice-cold 1× PBS to remove residual formaldehyde and glycine, cells were incubated in blocking buffer (final concentrations: 10% normal goat serum, 5% BSA, 0.1% Triton X-100, and 1× protease inhibitor cocktail) for 30 min. Cells were then double-labeled with an anti-Arc antibody (Bioss; 1:20,000 dilution per million cells) and an anti-NeuN antibody (Abcam; 1:20,000 dilution per million cells), along with DAPI (Thermo Fisher; 1:2,000 dilution). Cells were washed twice with PBPT buffer and resuspended in 500 μl of 1× PBS for FACS sorting using a BD FACSAria (BD Biosciences). Sorted cells were collected in 100 μl of PBS.

### Modified ecDNA purification from neuronal samples

Briefly, cells obtained from sorted cells or the mPFC of adult mice were used as input for nuclei isolation (Sigma). Alkaline-mediated chromosomal DNA precipitation was performed on the isolated nuclei samples (MACHEREY-NAGEL). Phenol-chloroform-isoamyl alcohol mediated DNA purification was performed after centrifugation to purify the crude ecDNA samples from the supernatant after centrifugation.

CTAB-mediated genomic DNA precipitation was performed after the crude ecDNA samples was obtained. 2% (w/v) of Cetyltrimethylammonium bromide (CTAB) solution in 40mM NaCl and 0.1M MgCl_2_ and was added to adjust the final concentration as 0.15%. After centrifugation at 14000rpm for 10 minutes, 3 vol. of absolute Ethanol, 0.1 vol. of 3M sodium acetate was added to the supernatant. After incubation at −20°C overnight, centrifugation at 14000rpm for 30 minutes was performed accompanied with 2 times of 80% ethanol wash. The pellet containing purified ecDNA was eluted in ultrapure water (Thermo Fisher).

Exonuclease-V mediated chromosomal DNA digestion was performed with 1ug of DNA obtained from the last step. 3ul of Exonuclease V (NEB), 5ul of buffer 4, 5ul of ATP, 5ul of Ampicillin and 5μl of 50% PEG-8000 (W/V) was added and incubated at 37°C with gentle mixing. A 7-day incubation with 1ul of ATP, 0.5ul of Exonuclease V and 0.5ul of buffer 4 were added every day. After the 7-day digestion, the exonuclease V was deactivated at 75°C for 10 minutes and 0.2 volume of AMPure XP beads was added. After incubation at ambient temperature for 5 minutes, the samples were placed in a magnetic rack and the supernatant was used as input for Phenol-chloroform-isoamyl alcohol mediated DNA purification. The purified DNA was quantified using a Qubit spectrometer and analysed with a Bioanalyzer 2100 and 1% agarose gel electrophoresis stained with 1% SYBR gold staining solution.

A quality check was performed for each step of ecDNA purification. *B2m* and *Pthlh* were used as reference genes to quantify the copy number of mtDNA through qPCR. In Brief, 1μl of 10uM primer pair, 5μl of 2x SensiFast SYBR NO ROX master mix (Meridian), 1μl of DNA template and 1μl of ultrapure water were mixed and qPCR reactions were carried out as follow: 95°C × 10’ for one cycle; 95°C × 10”, 60°C × 15”, 72°C× 20”, followed by plate read, for 50 cycles. Each PCR reaction was run in at least two technical replicates.

### DNA extraction

Samples were mixed with an equal volume of DNA extraction buffer (final concentrations: 0.1 M Tris-HCl, pH 7.5; 0.05 M EDTA, pH 8.0; 1.25% SDS; 0.002 mg/mL RNase A). Proteinase K (2 μl) was added, and the samples were incubated at 56 °C overnight. An equal volume of phenol:chloroform:isoamyl alcohol (Sigma; 25:24:1, v/v/v) was added, and the mixture was centrifuged at 14,000 rpm for 10 min at room temperature. The aqueous phase was collected and mixed with 3 vol. of absolute ethanol and 0.1 vol. of 3 M sodium acetate. Following overnight incubation at −20 °C, the mixture was centrifuged at 14,000 rpm for 30 min, and the resulting pellet was washed twice with 80% ethanol. The purified ecDNA pellet was then eluted in ultrapure water (Thermo Fisher).

### Super-resolution imaging

Purified ecDNA and corresponding genomic DNA samples were used for visualization of circular DNA molecules. Briefly, a 1 mM YOYO-1 stock solution (Invitrogen, Y3601) was diluted to a final concentration of 0.1 μM in 10 mM sodium phosphate buffer (1.88mM NaH_2_PO_4_H_2_O, pH adjusted to 7.5 with NaOH). For each 1 ng of DNA, 3 μl of the diluted YOYO-1 solution was added, and the mixtures were incubated at room temperature (~ 22 °C) for 2 h to ensure sufficient staining. Samples were imaged using a Zeiss Elyra 7 structured illumination microscope (SIM) with the following parameters: 488 nm high-resolution (HR) 200 mW diode laser (10–15% laser power), BP 495–575 nm/LP 750 nm filter set, and a Plan-Apochromat 63×/1.40 oil immersion objective (M27).

### Short-read low-input ecDNA sequencing

For each ecDNA sample, 1.5 ng of purified ecDNA was used as input for the Nextera XT DNA Library Preparation Kit (Illumina) with minor modifications to the standard protocol. Briefly, for each library, 10 μl of TD buffer and 3 μl of ATM buffer were mixed with the DNA sample. Tagmentation was performed by incubating the DNA-Tn5 transposome mixture at 37 °C for 32 min. The reaction was then neutralized by adding 5 μl of NT buffer and incubating at room temperature (approximately 22 °C). Library amplification was performed by adding 15 μl of NPM buffer and 10 μl of a Nextera UD index primer pair, followed by PCR using the following cycling conditions: initial denaturation at 95 °C for 30 s, followed by 15 cycles of denaturation at 95 °C for 10 s, annealing at 55 °C for 30 s, and extension at 72 °C for 30 s, with a final extension at 72 °C for 5 min. Following amplification, libraries were size selected using 0.6× volume of AMPure XP beads (Beckman Coulter). After a 5-min incubation at room temperature (approximately 22 °C), beads were washed twice with 80% ethanol. The purified and size-selected libraries were quantified and sequenced on an Illumina NovaSeq 6000 platform using 150-bp paired-end sequencing at GENEWIZ, Azenta Life Sciences.

### Full-length ecDNA sequencing

For full-length sequencing, the DNA input for library construction was prepared with rolling circle amplification (RCA) and debranching. RCA and debranching was performed with 100pg of ecDNA as template by Repli-G mini kit (Qiagen) according to the manufacturer’s instructions with minor modifications. The incubation was performed at 30°C for 48 hours. The purified DNA samples after 0.5 x volume AMPure XP bead selection was used as input for debranching. For each 20μl reaction mixtures, 2μl of 10 x phi29 DNA polymerase buffer, 5ul of phi29 DNA polymerase, 2μl of 25mM dNTP, and the purified DNA are added, and the incubation is performed at 30°C for 2 hours (NEB). After denaturation of phi29 DNA polymerase and phenol-chloroform-isoamyl alcohol mediated DNA purification, the further DNA debranching, mismatch depletion and repair was performed following the protocol described in this paper and the final samples were used for sequencing library construction with Sequel II RT sequencing kit (Pacific Biosciences, 101-826-100)^17^.

### ecDNA sequencing data analyses

#### Mapping of short read sequencing reads and ecDNA detection

Raw Illumina sequencing reads were trimmed for adapter removal and quality filtered using Trim Galore (v0.6.10) with default parameters. Trimmed reads were aligned to the mm10 mouse reference genome using BWA-MEM (v0.7.18) with default parameters. Circular DNA was detected using Circle-Map Realign. ecDNAs with a combined score >1 were selected for downstream analysis. Sequencing coverage was calculated using deepTools bamCoverage (v3.5.6). ecDNA coordinates were annotated using HOMER (v4.11), and GO analysis was performed using ShinyGO (v0.80).

#### Long-read base calling and read mapping

Consensus read BAM files were converted to FASTQ format using pbtk (v3.1.1) with default parameters. These FASTQ reads were aligned to the mm10 mouse reference genome using minimap2 (v2.28) with the -ax map-pb preset. ecDNA detection from the aligned reads was performed using FLED and ecDetector with the following setting ‘-j 25 -b 8 -S 2 -I 1 -D 1’ (v1.0.0). To validate ecDNAs identified from short-read sequencing, reads spanning the predicted ecDNA coordinates were extracted from the FASTQ files using bbmap filter (v39.13; rcomp=f mm=f hdist=1). These filtered reads were then aligned to the mm10 reference genome using minimap2 (v2.28) with the same parameters.

### Divergent PCR and sanger sequencing validation

PCR reactions were carried out in 10 μl volumes, comprising 1 μl of a 10 μM primer pair, 5 μl of EconoTaq PLUS GREEN 2X Master Mix (BioSearch Technologies), 1 μl of ecDNA-enriched template, and 1 μl of ultrapure water. The thermal cycling conditions included an initial denaturation step at 95 °C for 5 minutes, followed by 50 cycles of denaturation at 95 °C for 10 seconds, touchdown annealing from 70 °C to 58 °C (decreasing by 1 °C per cycle for the first 12 cycles, then maintaining 58 °C for the remaining cycles), and extension at 72 °C for 20 seconds. Each reaction was performed in at least two technical replicates.

Following amplification, PCR products were mixed with 6× DNA loading dye (NEB) and resolved via 2% agarose gel electrophoresis at 10 V/cm. DNA bands corresponding to the expected size were excised and purified using the Zymoclean Gel DNA Recovery Kit (Zymo), according to the manufacturer’s protocol. Purified amplicons (200 ng) were combined with 5 pg of either forward or reverse primers and submitted to GRS Sequencing Service at the University of Queensland for sequencing.

### RT-qPCR and qPCR quantification

qPCR reactions were performed in 10 μl volumes, consisting of 1 μl of a 10 μM primer pair, 5 μl of SensiFAST SYBR No-ROX Kit (Meridian), 1 μl of total DNA template, and 1 μl of ultrapure water. Thermal cycling conditions included an initial denaturation at 95 °C for 10 minutes, followed by 50 cycles of denaturation at 95 °C for 10 seconds, touchdown annealing from 70 °C to 58 °C (decreasing by 1 °C per cycle for the first 12 cycles and holding at 58 °C for subsequent cycles), and extension at 72 °C for 15 seconds. Each reaction was also performed in at least two technical replicates. For RT-qPCR, RNA isolation was performed using NucleoZOL (MACHEREY-NAGEL) following the protocol provided by the manufacturer. Followed by the reverse transcription using SensiFAST™ cDNA Synthesis Kit (Meridian) with the protocol provided by the manufacturer, the synthesized cDNA samples were diluted to approximately 100 ng/μl and used as template for qPCR reactions.

### ChIP-qPCR

ChIP was performed with modifications to the Invitrogen ChIP kit protocol. Primary cortical neurons were cross-linked with 1% formaldehyde in PBS. Cross-linked chromatin was sheared using a Covaris sonicator in 1% SDS lysis buffer to produce DNA fragments averaging 500 bp (sonication settings: peak power 75, duty factor 10%, cycles per burst 200, duration 900 s, temperature maintained at 5–9 °C). Chromatin was immunoprecipitated overnight at 4 °C with validated antibodies against TOP2B (Abcam) and Normal rabbit IgG (Cell Signaling Technology) served as a negative control. Antibody-chromatin complexes were captured using Protein G magnetic beads (Invitrogen) for 1 hour at 4 °C, followed by three washes each with high-salt and low-salt wash buffers. Proteinase K digestion was then performed on the washed beads, and DNA was purified via phenol-chloroform extraction and ethanol precipitation. The purified DNA was analysed by qPCR using primers targeting 200-bp regions of interest. Samples with insufficient enrichment relative to the IgG control were excluded from further analysis.

### DIP-seq Library Preparation and Sequencing

KCl-induced and naïve DIV-14 Primary cortical neurons were crosslinked with 2% paraformaldehyde solution for 10 minutes, quenched with 0.5M glycine solution for 5 minutes at room temperature and then incubated for 1 hour at 4°C in NP40 cell lysis buffer (Thermo Fisher) supplemented with protease inhibitor cocktail, RNaseOut and DTT, then sonicated with peak power as 75W; Duty Factor as 20% and 200 cycles per burst for 15 minutes. Sonicated samples were centrifuged at 14000rpm for 10 minutes at 4°C and the supernatant was transferred to a new tube, and the total amount of protein input was determined by Bradford Protein Assay. The protein lysate was diluted with dilution buffer (612.5mM NaCl; 6.25mM MgCl_2_) at a 1:4 ratio and pre-cleared with 5μl of Dynabeads MyOne Streptavidin T1 magnetic beads (Thermo Fisher). *In vitro* synthesis of sgRNA was performed using T7 RNA polymerase following the manufacture’s protocol (NEB).

For every 1mg of protein lysate, 1 μg of dCas9-FLAG-biotin was used, incubated with 3μg of in vitro transcribed guide RNA, was added to the lysate. After overnight incubation at room temperature with rotation, 20 μl of Streptavidin T1 beads was added and the slurry was incubated at room temperature for 2 hours. Beads, separated by the magnetic rack, was washed with Cell lysis buffer II supplied with DTT, protease inhibitor three times and the beads were used for library preparation and mass spectroscopy.

For DIP-seq, DNA was eluted from the beads by incubation with 200 μl of DNA extraction buffer (0.1 M Tris-HCl, pH 7.5; 0.05 M EDTA, pH 8.0; 1.25% SDS) containing 2 μl of proteinase K (NEB) overnight at 56 °C. Cross-linking was reversed, and DNA was purified as described above. Purified DNA was quantified using a Qubit dsDNA High Sensitivity assay. Libraries were prepared using the Nextera XT DNA Sample Preparation Kit (Illumina) according to the manufacturer’s protocol and sequenced on a NovaSeq 6000 platform (150-bp paired-end sequencing) at GENEWIZ, Azenta Life Sciences.

### DIP-Seq data analysis and Overlapping analysis

Cutadapt (v4.9) was used to clip low-quality nucleotides and adaptor sequences. Bases with a Phred quality score below 20 at the 3′ end were trimmed from each read. Subsequently, known Illumina primer and adaptor sequences were removed from each read using Cutadapt, which performs sensitive semi-global alignments allowing for gapped and mismatched alignments. BWA (v0.7.18) was used to align the clean reads against the mouse genome reference (mm10). SAM to BAM conversion, sorting, fixmate application, and removal of duplicate paired-end reads were all performed using Samtools (v1.8). For peak calling, MACS3 (v3.0.1) was used with the options “-f BAMPE -g mm -B -p 0.05” and the Narrowpeak files were used as input for consensus peak identification using MSPC with the parameters set as “-w 1e-4 -s 1e-8 -c 3”. For differentiated peak analysis, PePr with parameters “-- threshold 1e-4 --peaktype sharp --normalization intra-group --keep-max-duplicate 1” was set to confirm the differentiated binding peak regions. The occupancy of *ecCldn34d* was validated by DIP-qPCR using primers targeting 200-bp regions of interest.

### IP-MS Analysis

Mass spectrometry analysis was performed at the Mass Spectrometry Facility, Institute for Molecular Bioscience (IMB), University of Queensland. For protein digestion, magnetic affinity beads were incubated overnight at 37°C with 40 µL of sequence-grade trypsin (40 ng/µL in 50 mM ammonium bicarbonate buffer, pH 8; Promega). Following incubation, the trypsin solution was collected and transferred to fresh microcentrifuge tubes. The beads were then treated with 200 µL of 5% formic acid/acetonitrile solution (3:1, v/v) and incubated with shaking at room temperature for 30 minutes.

Prior to MS analysis, dried samples were reconstituted in 15 µL of 1.0% (v/v) trifluoroacetic acid (TFA). Tryptic peptides were analyzed using a microflow HPLC-MS/MS system consisting of an Eksigent Ekspert nanoLC 400 uHPLC coupled to a Triple TOF 6600 mass spectrometer (both from SCIEX) equipped with a micro Duo IonSpray ion source. Full-scan TOF-MS data were acquired over m/z 350-2000 with a 250 ms accumulation time, followed by up to 30 product ion (MS/MS) scans (50 ms each) using rolling collision energy in Information Dependent Acquisition (IDA) mode.

Data acquisition and processing were performed using Analyst software (version 1.7, SCIEX). Protein identification was conducted through database searching using ProteinPilot software (version 5.0, SCIEX).

### Synaptosome isolation and nuclear isolation

Synaptosomes and nuclei were isolated from primary cortical neurons and ILMPFCs. First, tissue was homogenized into a single-cell suspension using a Dounce tissue grinder. Synaptosomes were then isolated from this suspension using Syn-PER™ Synaptic Protein Extraction Reagent (Thermo Fisher Scientific) following the manufacturer’s protocol. From the remaining pellet separated during synaptosome isolation, nuclei were isolated using the Nuclei EZ Prep kit (Sigma-Aldrich) according to the manufacturer’s instructions.

### Molecular cloning

Lentiviral plasmids were generated by inserting ecCldn34d sgRNA or scrambled control fragments immediately downstream of the human EF1α promoter in the lentiCRISPR v2 vector (Addgene #52961) as previously described^18^.

### Lentiviral production and lentivirus titration

Lentivirus was prepared and maintained according to protocols approved by the University of Queensland. Either *ecCldn34d* knockout plasmid or the scramble control plasmid was co-transfected with lentiviral helper plasmids (pMDL, pVSVG and pREV) at a 2:1:1:1 ratio into HEK 293 T cells with roughly 80% confluence. Then 4 hours later, DMEM added with 10 mM of sodium butyrate was used to replace the media to stimulate the viral production. After 2 days’ incubation at 37 °C and 5% CO_2_, media containing the lentivirus was collected and ultra-centrifuged for harvesting. Titration of lentivirus was confirmed by real-time PCR after lentiviral transduction into HEK 293 T cells according to our previous publication^4^. Only viruses that reached over 1 × 10^8^ TU/ml were used in this study.

### Stereotaxic viral injection

Following a one-week habituation period, mice were anesthetized and bilaterally injected with lentivirus into the infralimbic prefrontal cortex (ILPFC). Using a 2 μL Hamilton syringe, a total of 1 μL per hemisphere was delivered at the following stereotaxic coordinates relative to bregma: AP +1.7 mm, ML ±0.2 mm, and DV −2.85 mm.

To minimize tissue damage, each injection was performed over a 20-minute period, consisting of a 10-minute infusion at 10 nl/min, followed by a 10-minute diffusion period where the syringe remained in place. After the injections, mice were recovered on a heating pad and monitored daily. Animals were allowed a recovery period of at least one week before subsequent experimentation.

### Behavioural tests

Behavioural training and testing were conducted in standard conditioning chambers (Coulbourn Instruments) featuring two transparent and two stainless-steel walls, with a steel grid floor for shock delivery (3.2 mm diameter rods, 8 mm apart). Animal behaviour was recorded by a ceiling-mounted digital camera, and freezing was quantified automatically using FreezeFrame software.

For fear conditioning, mice were placed in the experimental chamber (Context A), which was scented with a solution of 5% lemon and 10% alcohol. The conditioning protocol began with a 120-second baseline period, followed by three conditioning trials. Each trial consisted of a 120 s, 80 dB white noise conditioned stimulus (CS) that co-terminated with a 1 s, 0.7 mA foot shock unconditioned stimulus (US). A 120-second intertrial interval separated each CS-US pairing.

Fear memory retrieval was tested by returning the animals to Context A and presenting three CS-only trials (120 s white noise), each separated by a 120 s interval. The timing of this test varied between experimental groups: For animals conditioned before surgery, memory was tested 24 hours after conditioning. On the other hand, for animals conditioned after surgery, memory was tested 7 days after the microinjection procedure.

Freezing percentages were calculated for each trial and used for statistical analysis. Upon completion of the behavioural experiments, brains were processed to assess viral spread and knockout efficiency via immunohistochemistry according to a previously published protocol^18^ and qPCR.

## Acknowledgments

We gratefully acknowledge grant support from the NHMRC (GNT2003414). S.U.M. and E.L.Z. were supported by a Westpac Future Scholarship. We thank Dr. Alun Jones from the Mass Spectrometry Facility in the Institute of Molecular Bioscience at the University of Queensland for assistance with the proteomics experiments, the Queensland Brain Institute Advanced Microscopy Facility for its support.

## Figure legend

**Supplementary Fig. 1**. A. Summary of long-read sequencing metrics and the number of ecDNAs identified by ecDetector and FLED. B-C. Size distribution of ecDNAs identified in FACS-sorted NeuN+ (**b**) and NeuN-(**c**) cell populations (unpaired, two-sided Wilcoxon test; \*\*\*\**p* < 0.0001). Dashed lines indicate the average size for each population. D-E. Genomic feature annotation of ecDNAs from NeuN+ (**d**) and NeuN-(**e**) populations. F-G. TOP 10 GO Biological Process pathway analysis of genes associated with ecDNAs from NeuN+ (**f**) and NeuN-(**g**) populations. Pathways unique to the NeuN+ population are highlighted in red.

**Supplementary Fig. 2**. A. Validation of representative ecDNAs by gel electrophoresis. PCR amplicons spanning the predicted back-splice junction of each ecDNA are indicated by red arrows. B. Schematic outlining the strategy for full-length Sanger sequencing of *ecCldn34d*. C. Global alignment of the Sanger-sequenced ecCldn34d contig to its genomic locus of origin detected by Circle-Map and ecDetector. The junction site of *ecCldn34d* is indicated by red arrows, and sequence differences (polymorphisms and a microdeletion) are highlighted in red. D. Relative abundance of *ecCldn34d* in nuclear versus synaptosome fractions, as quantified by qPCR (n = 3 biologically independent samples; paired t-test; \*\**p* = 0.0083).

**Supplementary Fig. 3: Validation of *ecCldn34d* genomic interactome in neurons**. A, B, Genomic feature annotation of consensus *ecCldn34d* binding sites in neurons under basal (KCl−; a) and stimulated (KCl+; b) conditions. C, DIP-qPCR validation of activity-dependent *ecCldn34d* binding to putative sites on selected synapse-associated genes. (n ≥ 3 biologically independent samples, paired t-test; For pairwise comparison: \*\**p* < 0.01, \**p* < 0.05).

