## Supplementary Fig. 1 for "A mobile circular DNA element drives memory-related processes in mice"

**A**

| Sample | Subreads base (G) | Total reads | N50 Length (bp) | Longest Read (bp) | GC content (%) | ecDNA detected by ecdetector | ecDNA detected by FLED |
|---|---|---|---|---|---|---|---|
| NeuN+_rep1 | 3.3 | 681969 | 5714 | 36153 | 40 | 12153 | 2573 |
| NeuN+_rep2 | 4.3 | 822203 | 6252 | 36574 | 41 | | Overlapped: 2089 |
| NeuN-_rep1 | 4.1 | 777390 | 6289 | 47082 | 45 | 21074 | 10862 |
| NeuN-_rep2 | 3.9 | 682104 | 6855 | 43045 | 45 | | Overlapped: 10192 |

**B**

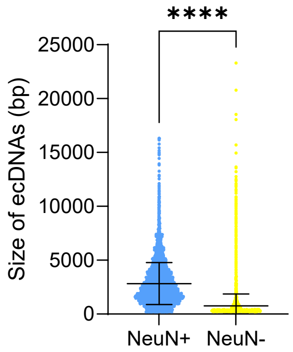


**C**

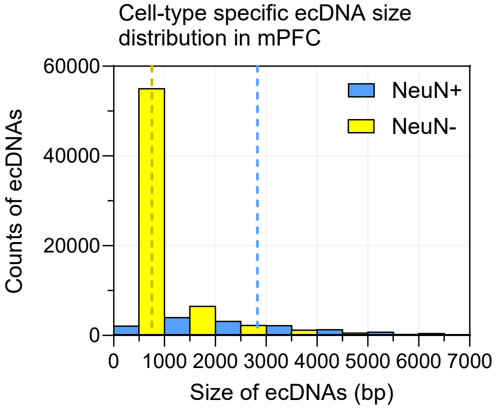


**D**

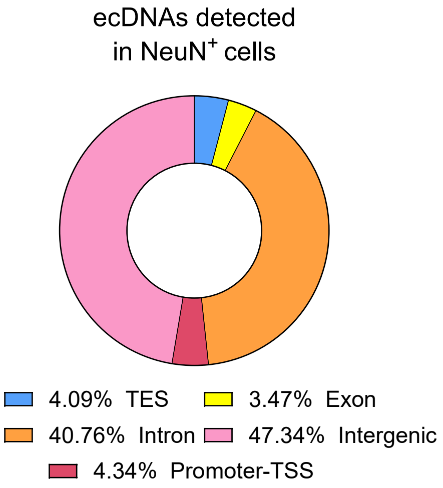


**E**

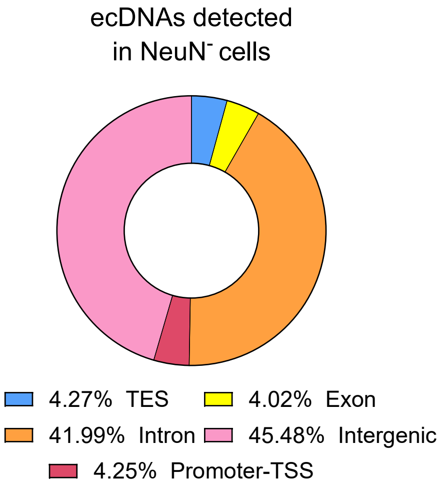


**F**

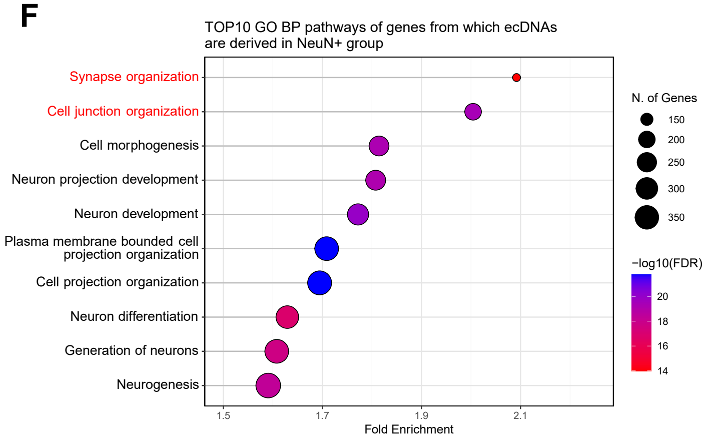


**G**

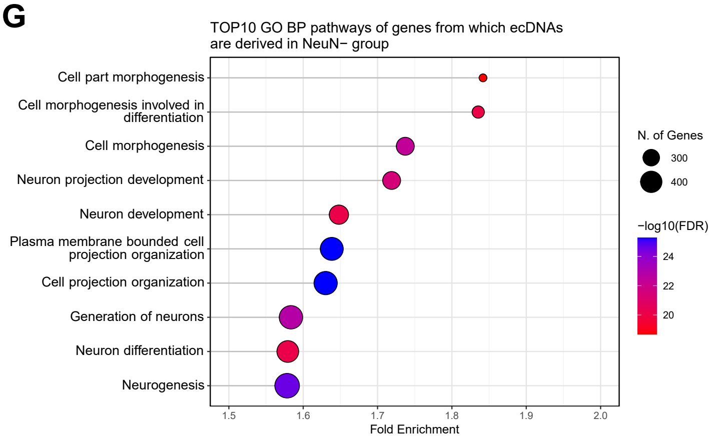


Figure S1
