## Supplementary Fig. 2 for "A mobile circular DNA element drives memory-related processes in mice"

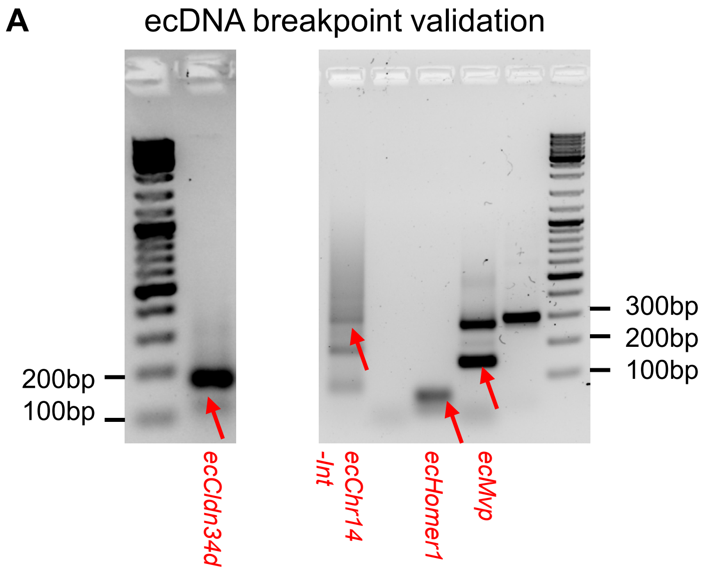


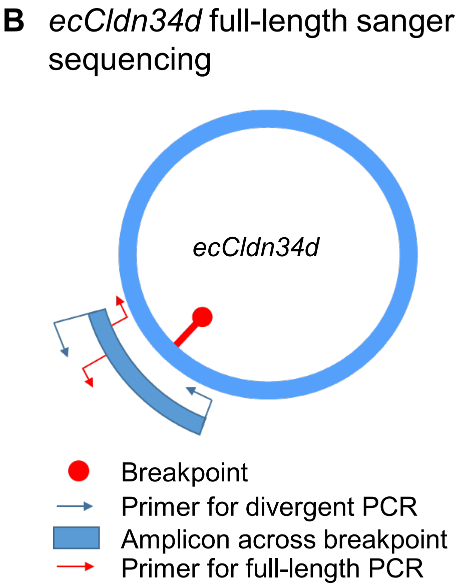


### C Alignment of *ecCldn34d* to the locus of origin

Aligned using Global alignment (Needleman-Wunsch)

```
ecCldn34d  ..  1  ACCTAGAAAATCAGACTGGCACACTGTAGAACATAGTGTGAAATGCACATTGTGAAACATAGTATATGAGTGTGTTGCACTGAAACATATAGAAAATTGGAAATTTCCAG  110
gCldn34d       1  ----AGAAAATCAGACCGGCACACTGTAGAACATAGCGTGAAATGCACATTGTAAAACATAGTATATGAGTGAGTTGCACCGAAACATATAGAAAATTTGAAATTCCCAG  106

ecCldn34d    111  TAGAGAACACAGTATATGAGTGAGTTAAACTAAAAAACCTAGAAAATAAGGCCGGCAAATAGTAGAACATAGTGTGAAATGCCCATTGT-AAACATAGTATGTTAATAAG  219
gCldn34d     107  TATAGAACACAGTATATGAGTGAGTTTCACTGAACAACCTAGAAAATAAGGCCTGCACACTGTAGAAAATAGTGTGAAATGCCCATTGTAAAACACAGTATGTTAATGAG  216

ecCldn34d    220  TTGCACGGAAAAACAGAAAATGGGAAATGCACAGTGTAGAACCCAATATATGAGTGAGTTACACTGAAACACCTAGAAAATCAGACCGGCACACTCTAGAACATAGTGTG  329
gCldn34d     217  TTGCACTGAAAAACAGAAAATGGAAAATGCAGAGTGTAGAAGCCAATATATGAGTGAGTTGCAATGAAACACCTTGAAAATGAGACCGCCACACTGTAGAACATAGTGTG  326

ecCldn34d    330  AAATGCATATTGTAAAACAAAGTATGTTTATTGAGTTGCACTG-AAAACAGAAAATGAGAAATGCACAGTGTAGAAAAAAAAATATATGGGTGAGTTGCAGTGAAACACC  438
gCldn34d     327  AAATGCACATTGTAAAACACAGTATG-TTAATGAGTTGCACTGAAAAACAGAAAATGGGAAATGCACAGTGTAG--AACCCAATATATGAGTGAGTTGCACTGAAACACC  433

ecCldn34d    439  TAGAAAATCAGACCGGCACACTGTAGAACATAGTGTGAAATGCACATTGTAAAACACAGGATGTTAATGAGTTGTGCTGAAAACCATAGAAAATGGGAAATGCACAGTGT  548
gCldn34d     434  TAGAAAATCAGACCGGCACATTGTAGAACATAGTGTGAAATGCACATTGTAAAACACAGGATGTTAATGAGTTGCACTGAAAAACATAGAAAATGGGAAATGCACAGTGT  543

ecCldn34d    549  AGAACATTGTATATGAGTGCTTTGCACT----------GAAAA  581  ..
gCldn34d     544  AGAACATTGTATATGAGTGCTTTGCACTGAACAACCTAGAAAA  586
```

Printed from SnapGene®

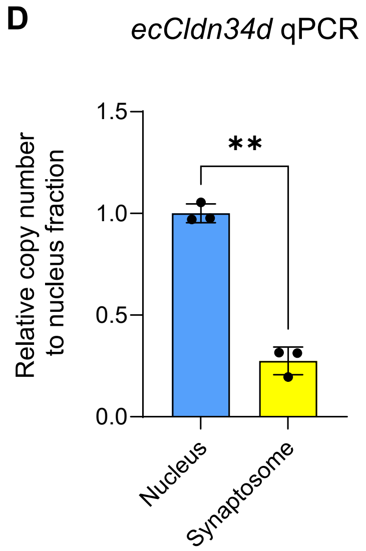


Figure S2
