## Supplementary figures and images for "A mobile circular DNA element drives memory-related processes in mice"

### Supplementary Fig. 3

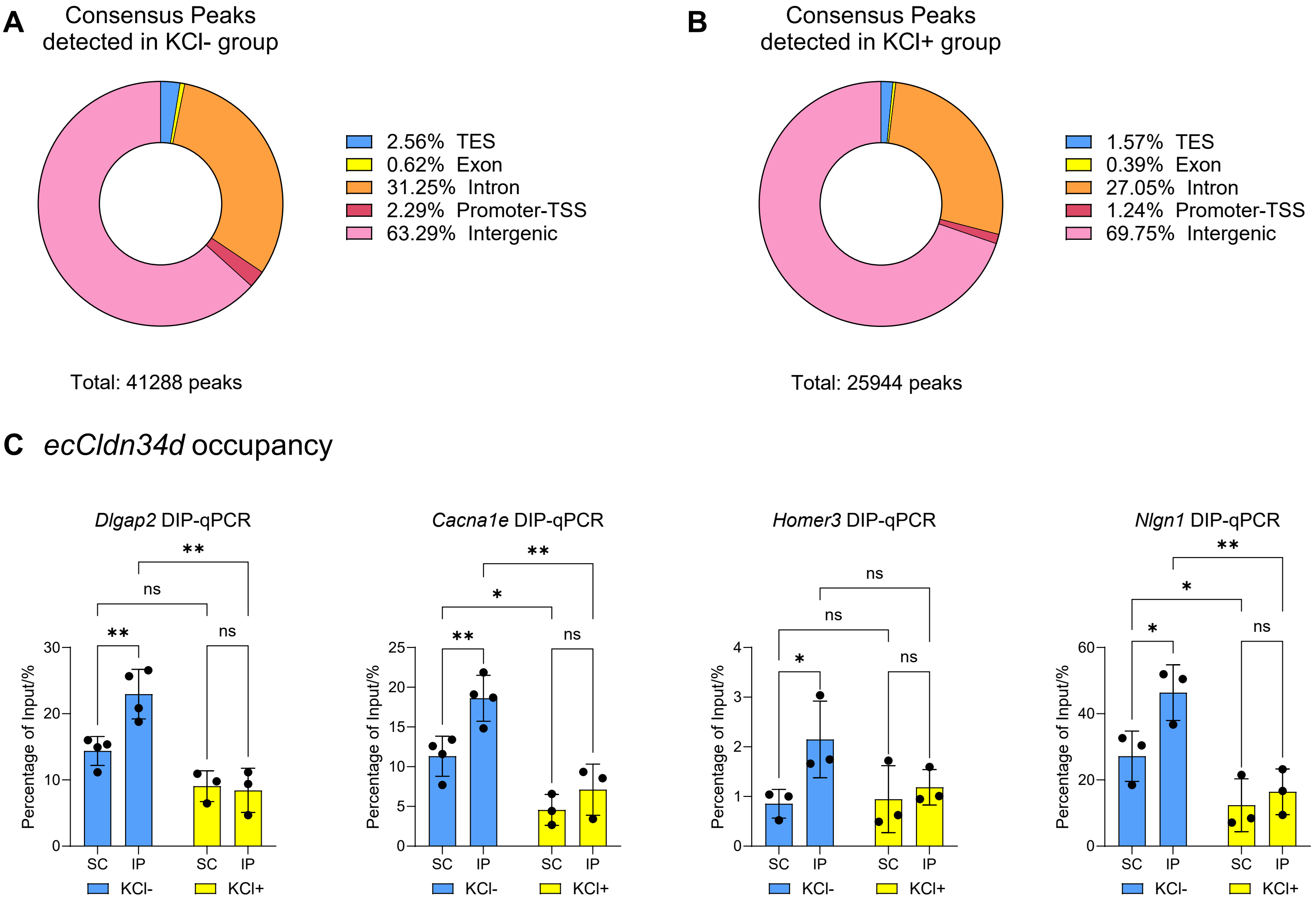
